# Development of a novel oral treatment that rescues gait ataxia and retinal degeneration in a phenotypic mouse model of familial dysautonomia

**DOI:** 10.1101/2022.11.04.515198

**Authors:** Elisabetta Morini, Anil Chekuri, Emily M. Logan, Jessica M. Bolduc, Emily G. Kirchner, Monica Salani, Aram J. Krauson, Jana Narasimhan, Vijayalakshmi Gabbeta, Shivani Grover, Amal Dakka, Anna Mollin, Stephen P. Jung, Xin Zhao, Nanjing Zhang, Sophie Zhang, Michael Arnold, Matthew G. Woll, Nikolai A. Naryshkin, Marla Weetall, Susan A. Slaugenhaupt

## Abstract

Familial Dysautonomia (FD) is a rare neurodegenerative disease caused by a splicing mutation in the Elongator complex protein 1 gene (*ELP1*). This mutation leads to the skipping of exon 20 and a tissue-specific reduction of ELP1 protein, mainly in the central and peripheral nervous systems. FD is a complex neurological disorder accompanied by severe gait ataxia and retinal degeneration. There is currently no effective treatment to restore ELP1 protein expression in individuals with FD, and the disease is ultimately fatal. After identifying kinetin as a small molecule able to correct the *ELP1* splicing defect, we worked on its optimization to generate novel splicing modulator compounds (SMCs) that can be used in patients. Here, we optimize the potency, efficacy, and bio-distribution of second-generation kinetin derivatives to develop an oral treatment for FD that can efficiently pass the blood-brain barrier and correct the *ELP1* splicing defect in the nervous system. We demonstrate that the novel compound, PTC258, efficiently restores correct *ELP1* splicing in mouse tissues, including brain, and most importantly, prevents the progressive neuronal degeneration that is characteristic of FD. Postnatal oral administration of PTC258 to the phenotypic mouse model *TgFD9;Elp1^Δ20/flox^* increases full-length *ELP1* transcript in a dose-dependent manner and leads to a two-fold increase in functional ELP1 protein in the brain. Remarkably, PTC258 treatment improves survival, gait ataxia, and retinal degeneration in the phenotypic FD mice. Our findings highlight the great therapeutic potential of this novel class of small molecules as an oral treatment for FD.

## Introduction

Familial dysautonomia (FD), also known as Riley-Day syndrome or hereditary sensory and autonomic neuropathy type III (OMIM223900, MIM603722), is an autosomal recessive neurodegenerative disease caused by a splicing mutation in the Elongator complex protein 1 (*ELP1*, formerly called *IKBKAP*). This mutation results in variable tissue-specific skipping of exon 20 with a corresponding reduction of ELP1 protein in the central and peripheral nervous systems. ELP1 is the scaffolding subunit of the human Elongator complex, a highly conserved protein complex that participates in distinct cellular processes, including transcriptional elongation, acetylation of cytoskeletal α-tubulin, and tRNA modification ^1–11^. ELP1 function has been extensively studied and has been implicated in exocytosis, cytoskeletal organization, axonal transport, and cellular adhesion and migration ^12–16^. Recent studies highlighted the role of ELP1 in neurogenesis, neuronal survival, and peripheral tissue innervation ^17–23^.

FD occurs almost exclusively in Ashkenazi Jews, with a carrier frequency of 1 in 32 in the general Ashkenazi Jewish population and 1 in 19 in Ashkenazi Jews of Polish descent ^24,25^. The clinical features of FD are all due to a striking progressive depletion of unmyelinated sensory and autonomic neurons^26–30^. Patients with FD have a complex neurological phenotype that includes diminished pain and temperature perception, decreased or absent myotatic reflexes, proprioceptive ataxia, and progressive retinal degeneration ^28,31–39^. The lack of afferent baroreceptor signaling causes complete failure of blood pressure regulation, and the recurrent hypertensive vomiting attacks are referred to as “dysautonomic crises” ^27,40–43^. Unexplained sudden death, aspiration pneumonia, and respiratory insufficiency remain the leading causes of death ^32,44^.

Many debilitating symptoms of the disease are due to progressive impairment of proprioception and retinal degeneration ^38,39,44–46^. Lack of afferent signaling from the muscle spindles accounts for the absence of deep tendon reflexes and gait ataxia ^39,45^. Children with FD are uncoordinated and tend to fall. As they age, progressive impairment in proprioception leads to severe gait ataxia, and they eventually lose the ability to ambulate independently ^39,45^. Neuropathological analysis of autopsy material from individuals with FD showed grossly reduced volume and number of neurons in the dorsal root ganglia (DRG) ^30^. Another debilitating aspect of FD that severely affects patients’ quality of life is progressive retinal degeneration, leading to visual dysfunction ^35,47,48^. Initially, it was reported that the loss of vision in FD patients resulted from corneal opacities, neovascularization, and sensory defects such as corneal analgesia, severe dry eye, ulceration healing, and incomplete closure of eyelids ^49–53^. However, recent detailed studies have shown that decreased visual acuity, loss of central vision, and temporal optic nerve pallor occur in FD patients even without any corneal complications, suggesting a neuro-ophthalmic nature of the disease ^47^. In FD, visual impairment is usually early-onset and often progresses to legal blindness in the third decade of life ^35^. Individuals with FD show a significant reduction in the retinal nerve fiber layer (RNFL) due to the death of retinal ganglion cells (RGCs) ^35,47,48^.

The field has undertaken many efforts to develop novel therapies to correct *ELP1* splicing defects, including splicing modulator compounds (SMCs), antisense oligonucleotides (ASO), and modified exon-specific U1 small nuclear RNAs (snRNAs) ^54–56^. Despite significant progress, we do not yet have a systemic therapy to prevent disease progression. Our team identified the small molecule kinetin (6-furfurylaminopurine) as an orally active splicing modulator of *ELP1* both *in vitro* and *in vivo* ^57,58^. As part of the NIH Blueprint Neurotherapeutics Network, we have improved kinetin potency and efficacy and generated a more potent and efficacious *ELP1* splicing modulator, BPN-15477 ^59,60^. More recently, our collaboration with PTC Therapeutics, Inc. led to the generation of a new class of highly potent SMCs, using BPN-15477 as a starting molecule, and to the identification of the novel compound PTC258. In this study, we describe the medicinal chemistry optimization of our new class of SMCs and we evaluate the efficacy of PTC258 in rescuing disease phenotypes in the FD mouse model *TgFD9; Elp1^Δ20/flox^*.

## Results

### Identification of the highly potent splicing modulator PTC258

As part of the NIH Blueprint Neurotherapeutics Network, we identified a class of SMCs that selectively modulate *ELP1* pre-mRNA splicing and increase the inclusion of exon 20 ^60,61^. Here, we optimized the potency, efficacy, and distribution of these compounds to develop an oral treatment for FD that could efficiently cross the blood-brain barrier (BBB) and correct *ELP1* splicing defect in PNS and CNS. In collaboration with PTC Therapeutics Inc., we have generated hundreds of novel BPN15477-analogs and identified PTC258 as a potent and specific *ELP1* splicing modulator. All components of the BPN15477 molecule were probed with systematic structural modifications. Right/Eastern substitution was most tolerated and provided the most substantial gains in potency. In particular, the primary amino group attached by a 2-carbon chain was optimal when substituted at the 2-position with small alkyl groups, as exemplified by compound PTC102, where a 30X boost in potency was observed (Figure 1A). The stereochemistry at the 2-position was found to be very important, favoring the (S)-enantiomer (for methyl-substitution). Additional optimization was achieved by replacing the pyrrolopyrimidine core with thienopyridine. Not only could an additional >30X improvement in potency be achieved over PTC102, but the thienopyridine analogs, including PTC258, showed superior oral exposure in mice (Figure 1A). PTC258 efficiently increases full-length *ELP1* mRNA and protein in FD patient fibroblasts (Figure 1B and C), and it is about 30,000 times more potent than kinetin (EC_2X_ ELP1 protein = 10,000 nM) and about 1,000 times more potent than BPN15477 (EC_2X_ ELP1 protein = 340 nM)^60^. We then assessed the ability of PTC258 to correct *ELP1* splicing and increase the amount of functional protein *in vivo*. We orally administered PTC258 to the *TgFD9* transgenic mouse ^62^, which carries the human *ELP1* gene with the major FD splice mutation, and mice were sacrificed on the 7^th^ day of treatment. Special chow was formulated to dose each mouse 3 mg/kg/day (0.002% PTC258 diet), 6 mg/kg/day (0.004% PTC258 diet), 12 mg/kg/day (0.008% PTC258 diet) and 24 mg/kg/day (0.016% PTC258 diet). While this mouse is phenotypically normal, as it expresses normal amounts of endogenous *Elp1*, it is a great model to initially assess the effect of SMCs on *ELP1* splicing *in vivo* because it recapitulates the same tissue-specific splicing defect observed in patients ^62,63^. PTC258 increased full-length *ELP1* transcript in a dose-dependent manner and, importantly, led to at least a 5-fold increase in functional ELP1 protein in the brain, trigeminal, liver, and quadricep (Fig. 1 D and E, Supplementary Fig. 1A and B). In addition, the treatment was well tolerated, no weight loss or adverse effects were observed in the treated groups, even at the highest concentration, and the level of splicing correction correlated with PTC258 exposure (Supplementary Fig.1C and D). These results demonstrate that treatment with PTC258 corrects splicing of the *ELP1* transcript and significantly increases the amount of functional protein *in vivo* in all tissues tested, including the brain.

**Figure 1.**
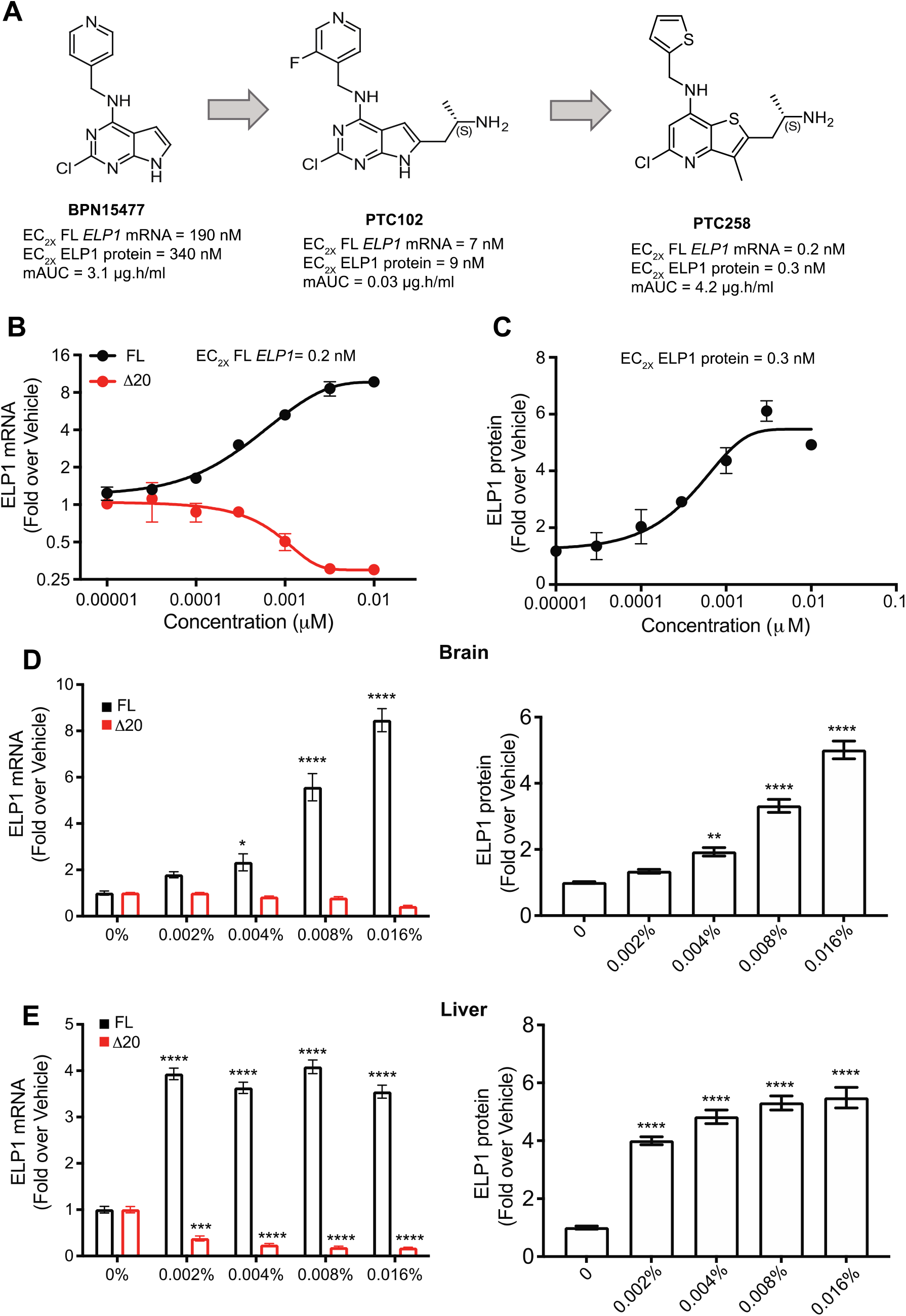
Identification of the novel small molecule splicing modulator PTC258. (**A**) Chemical optimization of BPN15477 to generate a more potent splicing modulator. Compounds were screened based on increasing the amount of full-length (FL) ELP1 mRNA, measured by qRT-PCR, and ELP1 protein, measured by Homogeneous Time Resolved Fluorescence (HTRF), in FD patient fibroblasts. EC_2X_ is the Effective concentration of the drug that achieves a 2-fold change in biological response relative to baseline. mAUC is the mouse area under the plasma drug concentration-time curve, reflects the actual body exposure to drug after administration of a dose of the drug and is expressed μg.h/ml. (**B**) Representative dose response curve of mutant (Δ20) and full-length (FL) *ELP1* transcripts in FD fibroblasts treated with increasing concentration of PTC258. Cells were treated for 24 h at the concentrations indicated (n = 6). (**C**) ELP1 protein expression in FD fibroblasts treated with increasing concentration of PTC258. Cells were treated for 24 h at the concentrations indicated (n = 6). (**D** and **E**) Relative expression of full-length (FL) and Δ20 ELP1 mRNA (left panel), and ELP1 protein quantification (right panel) in brain and liver after oral administration of PTC258 in chow ranging from 0.002% to 0.016% in adult transgenic *TgFD9* mouse (n =6-11). The adjusted P values are displayed. *P< 0.05, **P< 0.01, ***P< 0.001 and ****P< 0.0001, one-way ANOVA with Dunnett’s multiple comparison test. Data are shown as average ± s.e.m.

### Daily consumption of PTC258 improves gait ataxia and rescues retinal degeneration in the FD phenotypic mouse

To assess the therapeutic efficacy of PTC258 on disease progression, we administered the treatment to the phenotypic FD mouse model *TgFD9; Elp1^Δ20/flox^* ^64^. Special chow was formulated to dose each mouse 3 mg/kg/day (0.002% PTC258 diet) or 6 mg/kg/day (0.004% PTC258 diet). At birth, pups were randomly assigned to the vehicle or one of the two PTC258-treatment groups. Mice were maintained in the same treatment regime for the entire trial duration and were sacrificed at 6 months of age for tissue collection. Our previous studies in the FD mouse show that by 6 months of age the disease phenotype is evident and quantifiable ^56,59^. We started the treatment at birth to maximize the therapeutic value, and a preliminary study that assessed ELP1 expression in transgenic pups after PTC258 treatment showed that this compound can pass from dams to pups through lactation and increase the functional ELP1 protein amounts in the pups (Supplementary Fig. 2). The treatment was well tolerated as no weight loss was observed in the PTC258-treated group (Supplementary Fig. 3A and B). Notably, PTC258 improved the survival of FD pups in a dose-dependent manner (Supplementary Fig. 3C). Since our mouse model correctly recapitulates the gait ataxia observed in patients^39,45^, we first evaluated the effect of treatment on gait at three and six months of age ^56,64^. FD mouse gait was assessed using CatWalk XT (Noldus), a complete gait analysis system for quantitative assessment of footfalls and locomotion in mice ^65–67^. Particularly, stride length and base of support are two of the most relevant parameters for assessing gait in mice ^68^. The first is defined as the distance between successive placements of the same paw, while the latter represents the mean distance between hind paws (Fig. 2A). Our results show that the PTC258-treated FD mice exhibit a dose-dependent improvement in motor coordination at six months of age, as demonstrated by a progressive increase in stride length in PTC258-treated FD mice when compared with the vehicle-treated FD mice (Fig. 2B and C). The base of support was completely rescued in the 0.002% PTC258-treated group. In the 0.004% PTC258-treated group, although we observed a trend toward an increased base of support, the difference with the vehicle-treated FD mice was not statistically significant. One explanation can be that the sample size of the 0.004% PTC258-treated group was smaller compared with the 0.002% PTC258 treatment group. As expected by the progressive nature of the disease, at three months of age, FD mice do not yet exhibit dramatic gait abnormalities, except for the base of support that is already significantly lower in the vehicle-treated mice when compared with control or PTC-treated FD mice (Supplementary Fig. 4).

**Figure 2.**
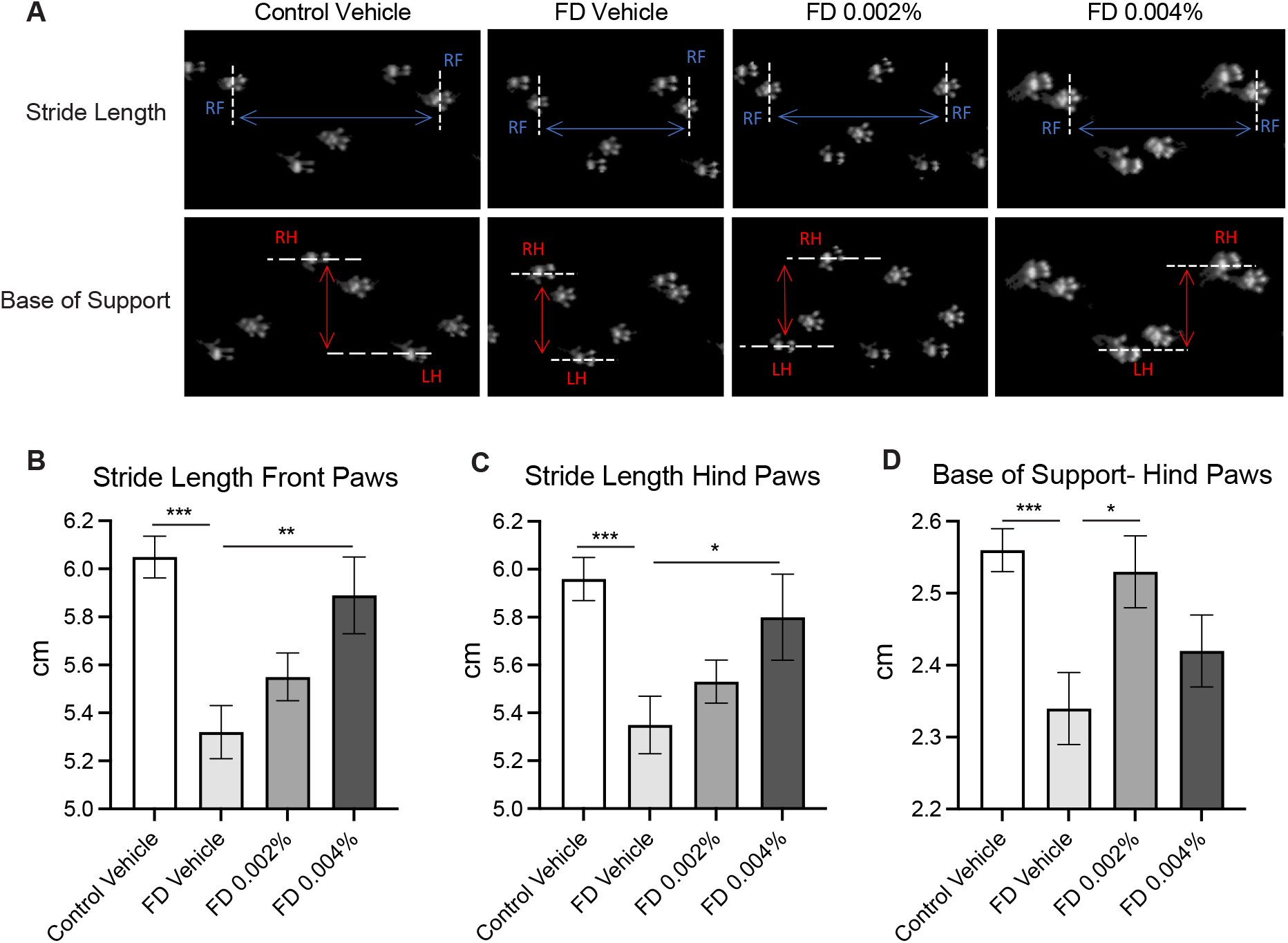
PTC258 improves motor coordination in the FD mice. (**A**) Representative Catwalk images of vehicle-treated control mice and vehicle-treated and PTC258-treated FD mice at 6 months of age. Stride length is defined as the distance between successive placements of the same paw (blue double-headed arrows), while the base of support represents the mean distance between hind paws (red double-headed arrows). Measurements of stride length front paws (**B**), stride length hind paws (**C**) and base of support hind paws (**D**) in vehicle-treated control mice (n=20) and vehicle-treated (n=16), 0.002% PTC258-treated (n=16) and 0.004% PTC258-treated (n=9) FD mice at 6 months of age. The adjusted P values are displayed. *P< 0.05, **P< 0.01 and ***P< 0.001, two-tailed unpaired Student’s t-test with FDR correction. Data are shown as average ± s.e.m.

Given that blindness is a debilitating aspect of FD, we evaluated for the first time the effect of our oral therapy to rescue retinal degeneration in the FD mice. Patients with FD show thinning of the RNFL layer due to the death of RGCs, and this loss is more profound near the temporal region of the optic nerve, specifically in the maculo-papillary region ^35,47,48^. High-definition spectral-domain optical coherence tomography (SD-OCT) was used to measure the thickness of the RFNL and the ganglion cell-inner plexiform layer (GCIPL) in the superior, inferior, nasal, and temporal hemispheres of the mouse retina (Fig. 3A) ^59^. At both ages, three and six months, we observed a significant reduction of the RNFL (Fig. 3B and D) and GCIPL (Fig. 3C and E) layers in each region of the vehicle-treated FD retinas. PTC258-treated FD mice showed a significant dose-dependent improvement in the thickness of both RNFL (Fig. 3B and D) and GCIPL (Fig. 3C and E). These results indicate that oral administration of PTC258 starting at birth prevents gait ataxia and retinal degeneration in the phenotypic FD mouse.

**Figure 3.**
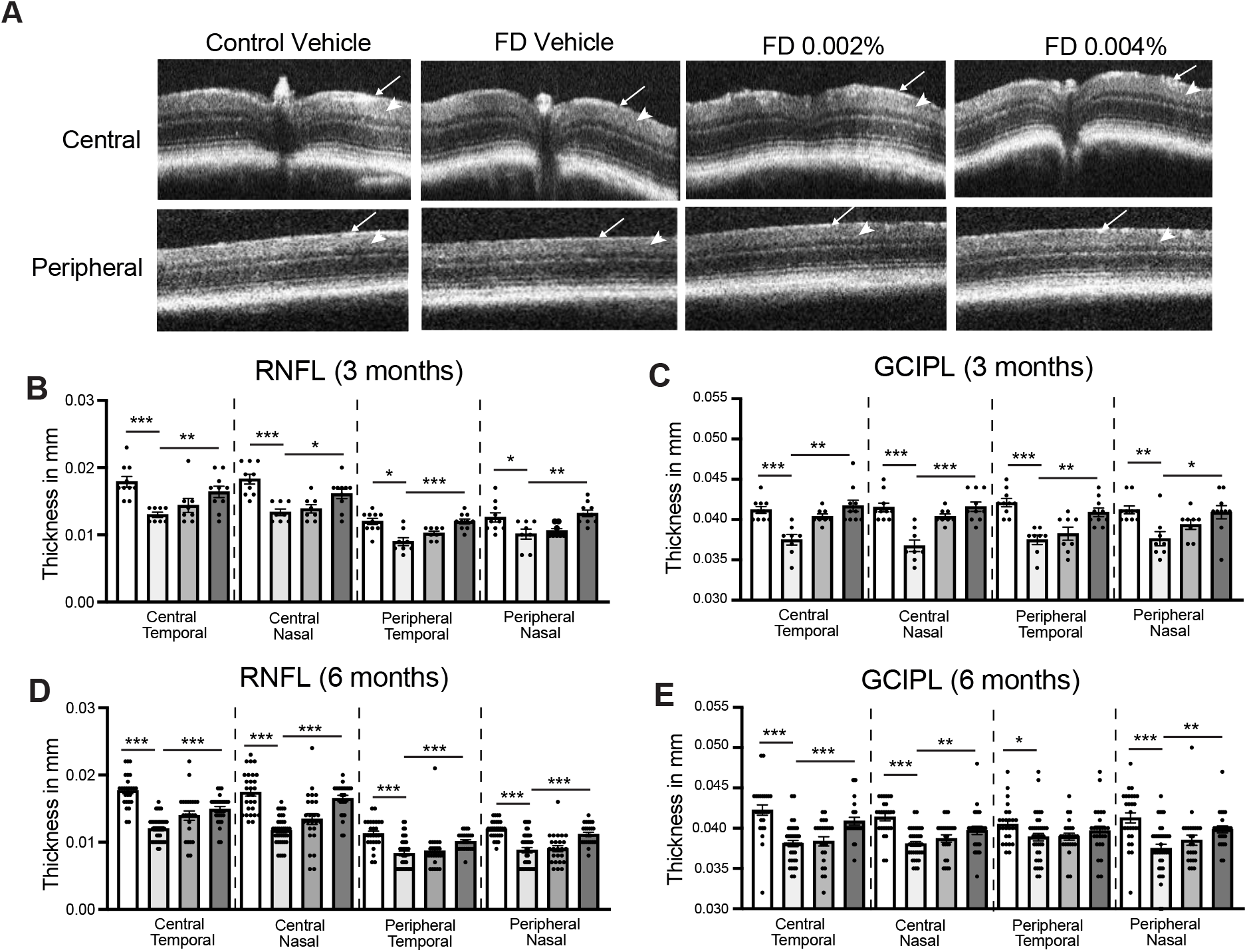
PTC258 rescues retinal degeneration in the FD mice. (**A**) Representative SD-OCT b-scan images of vehicle-treated control retinae and vehicle-treated and PTC258-treated FD retinae at 6 months of age. Arrows indicate the retinal nerve fiber layer (RNFL) while arrowheads indicate the ganglion cell-inner plexiform layer (GCIPL). Thickness measurements of RNFL (**B**) and GCIL (**C**) in both central and peripheral regions of the retina in nasal and temporal hemispheres at 3 months in vehicle-treated control retinae (n=10), and vehicle-treated (n=8), 0.002% PTC258-treated (n=8) and 0.004% PTC258-treated (n=10) FD retinae. (**D**) Thickness measurements of RNFL (**D**) and GCIL (**E**) in both central and peripheral regions of the retina in nasal and temporal hemispheres at 6 months in vehicle-treated control retinae (n=26), and vehicle-treated (n=38), 0.002% PTC258-treated (n=24) and 0.004% PTC258-treated (n=25-26) FD retinae. White bars represent control mice, light grey bars represent vehicle-treated FD mice, grey bars represent 0.002% PTC258-treated FD mice and dark grey bars represent 0.004% PTC258-treated FD mice. The adjusted P values are displayed. *P< 0.05, **P< 0.01 and ***P< 0.001, two-tailed unpaired Student’s t-test with FDR correction. Data are shown as average ± s.e.m., each data point represents an individual retina.

### PTC258 treatment prevents neuronal loss in FD DRG and retina

To confirm that the observed PTC258-mediated phenotypic improvement correlated with changes in the neuropathological hallmarks of the disease, we histologically characterized DRG and retinas from vehicle- and PTC258-treated mice. Individuals with FD have compromised fetal development and postnatal maintenance of DRG neurons, resulting in DRG of grossly reduced size and significantly reduced neuronal number ^30,69^. Proprioceptors are the subpopulation of neurons within the DRG responsible for sensory-motor coordination ^70,71^. Consistent with the observed proprioceptive deficits, vehicle-treated FD mice showed a significant reduction in the volume of the DRG, and the number of proprioceptive neurons compared with their control littermates (Fig. 4A). The DRG volume in the FD mouse was 60% of the controls (Fig. 4B), while the number of the proprioceptive neurons was reduced to 55% compared with their control littermates (Fig. 4C). Importantly, the treatment was able to rescue both neuropathological aspects of the disease. PTC258-treated FD mice showed a significant increase in the number of proprioceptive neurons and volume of DRG (Fig. 4 B and C), demonstrating that starting the treatment at birth is sufficient to prevent the loss of this subpopulation of neurons.

**Figure 4.**
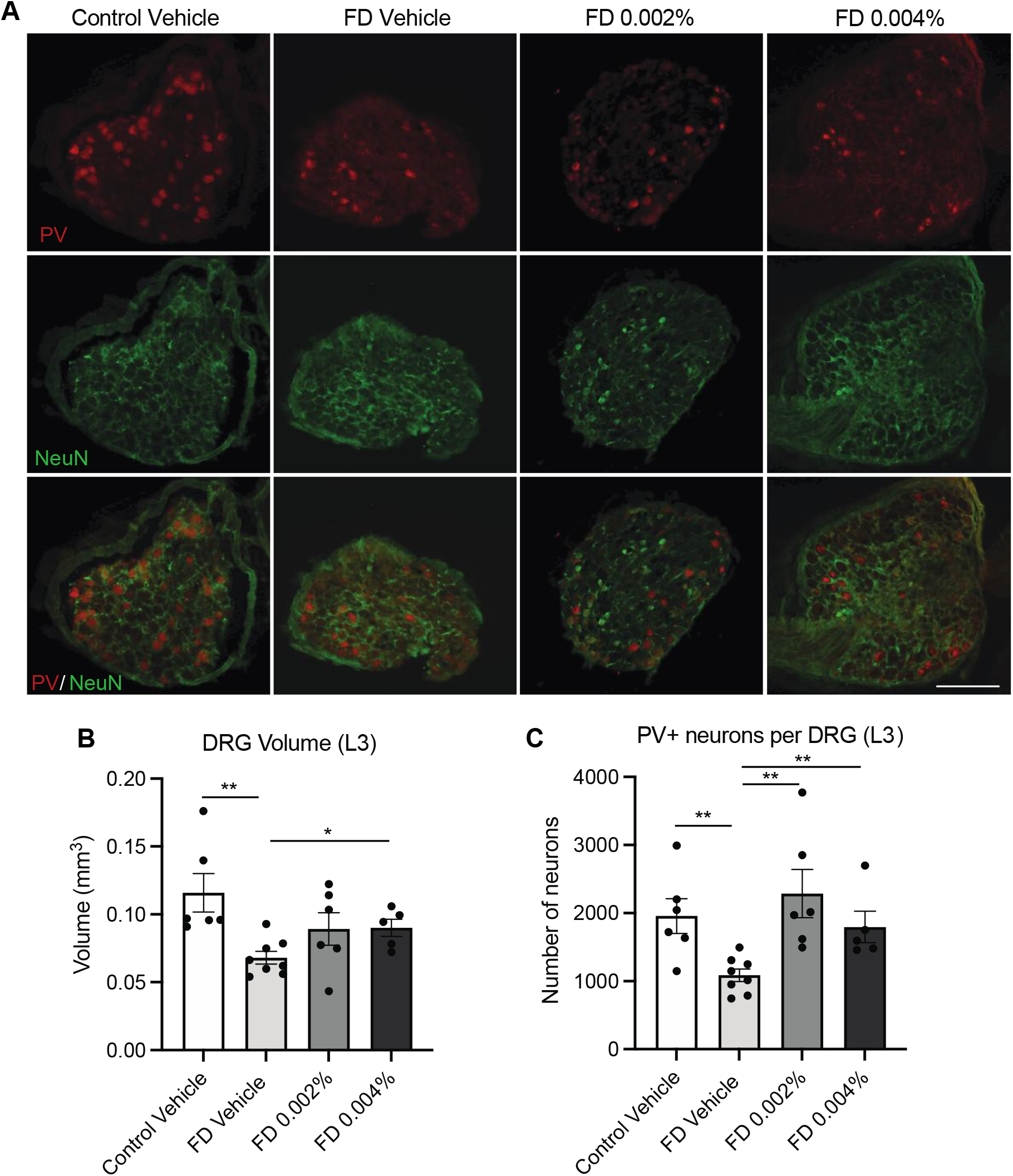
PTC258 treatment rescues proprioceptive sensory loss in the FD mice. (**A**) Representative images of proprioceptive (PV+) neurons (red), whole sensory (Nissl+) neurons (green) and the merged image (bottom) in L3 DRG from vehicle-treated control mice and vehicle-treated and PTC258-treated FD mice at 6 months of age. Scale bar, 200 μm. (**B**) Total volume of the L3 DRG measured in vehicle-treated control mice (n =5), and vehicle-treated (n=8), 0.002% PTC258-treated (n=6) and 0.004% PTC258-treated (n=5) FD mice. (**C**) Total number of PV+ proprioceptive neurons per DRG counted in vehicle-treated control mice (n =5), and vehicle-treated (n=8), 0.002% PTC258-treated (n=6) and 0.004% PTC258-treated (n=5) FD mice. The adjusted P values are displayed. *P< 0.05 and **P< 0.01, two-tailed unpaired Student’s t-test with FDR correction. Data are shown as average ± s.e.m., each data point represents an individual animal.

To evaluate whether the rescue of RNFL and GCIPL thickness was due to the treatment effect on RGC survival, we performed RGC counts in the superior, inferior, nasal, and temporal regions of the retina using retinal whole-mount analysis. We stained retinas from 3 and 6-month-old mice using the RGC-specific marker RNA-binding protein with multiple splicing (RPBMS), and we counted the number of RPBMS^+^ cells sited at 1mm from the optic nerve head (ONH) (Fig. 5A). As previously reported^59^, the FD mice at three months of age did not yet show a significant loss in RGC (Fig. 5B). However, in accordance with the progressive nature of retinal degeneration in FD patients, at six months of age, the number of RPBMS^+^ cells became significantly lower in the vehicle treated-FD mice in the temporal, nasal, superior and inferior regions of the retina (Fig. 5C). Importantly, PTC258 treatment rescued RGC loss in a dose-responsive manner (Fig. 5C). Together, these results demonstrate that our oral treatment improves disease phenotype by halting the loss of specific neuronal populations in FD DRG and retina.

**Figure 5.**
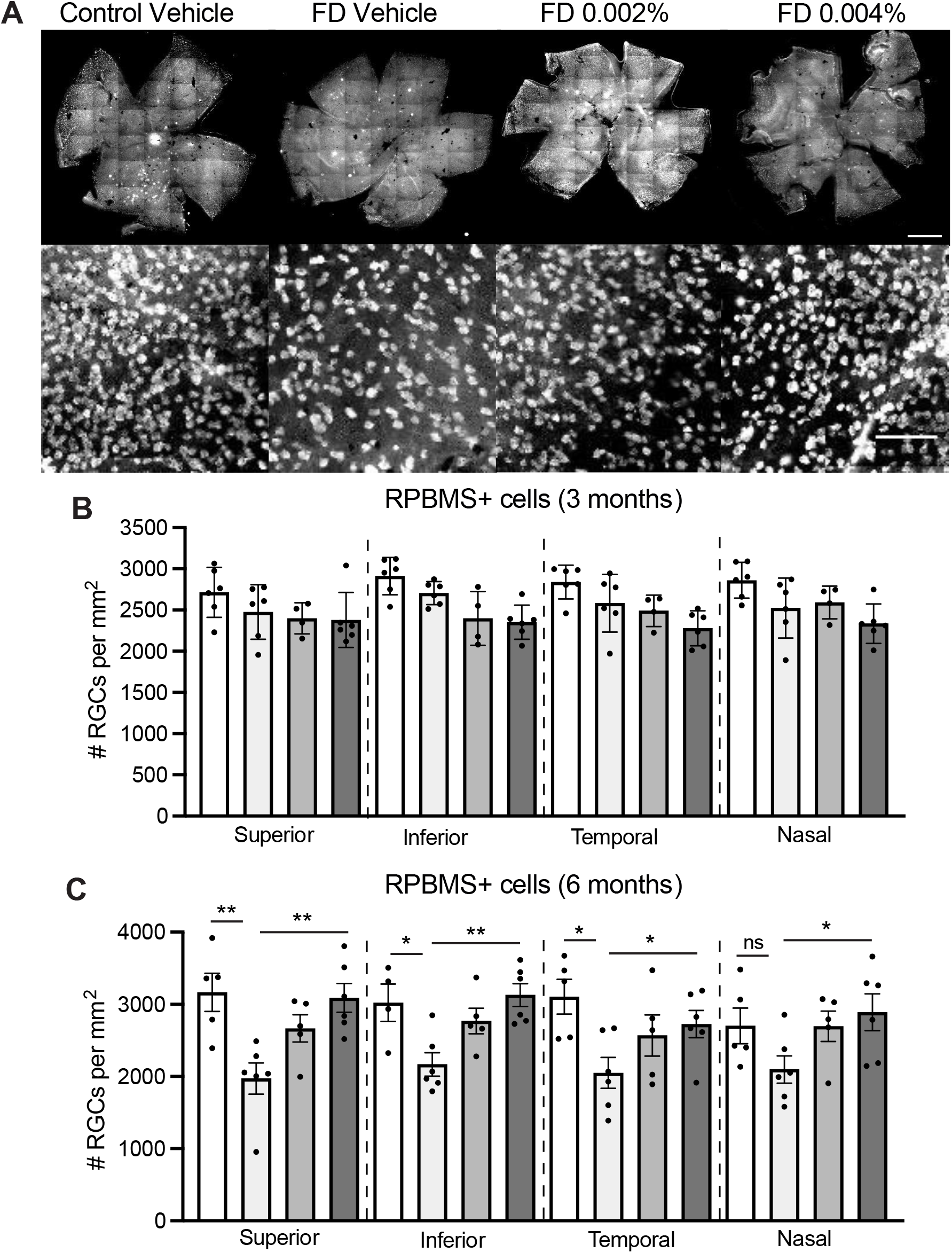
PTC258 prevents RGC loss in FD retinas. (**A**) Representative retinal whole-mount (top) and higher magnification of RPBMS staining (bottom) from vehicle-treated control mice and vehicle-treated and PTC258-treated FD mice stained with RGC marker RBPMS (top panels). Scale bars, 100 μm (top) and 0.5 μm (bottom). RPBMS+ cells were counted in each quadrant in superior (S), inferior (I), nasal (N), and temporal regions at 1mm from the optic nerve head (ONH) at 3 and 6 months of age. (**B**) Bar plots of RPBMS^+^ cell counts in 3 month-retinae from vehicle-treated control mice (n=6), and vehicle-treated (n=6), 0.002% PTC258-treated (n=4) and 0.004% PTC258-treated (n=6) FD mice. (**C**) Bar plots of RPBMS^+^ cell counts in 6 month-retinae from vehicle-treated control mice (n=5), and vehicle-treated (n=6), 0.002% PTC258-treated (n=5) and 0.004% PTC258-treated (n=6) FD mice. White bars represent control mice, light grey bars represent vehicle-treated FD mice, grey bars represent 0.002% PTC258-treated FD mice and dark grey bars represent 0.004% PTC258-treated FD mice. The adjusted P values are displayed. *P< 0.05 and **P< 0.01, two-tailed unpaired Student’s t-test with FDR correction. ns: not significant. Data are shown as average ± s.e.m., each data point represents an individual retina.

### PTC258 corrects *ELP1* splicing defect in PNS and CNS

Finally, we assessed if the phenotypic and neuropathological improvement observed in the treated FD mice correlates with the correction of the underlying FD splicing defect in PNS and CNS. *ELP1* splicing and protein amounts were analyzed in different neuronal tissues and liver from vehicle- and PTC258-treated FD mice. As shown in Figure 6, PTC258 treatment significantly increases *ELP1* exon 20 inclusion in the brain (Fig. 6A), DRG (Fig. 6B), trigeminal (Fig. 6C), retina (Fig. 6D), and liver (Fig. 6E). As expected, the improvement of exon 20 inclusion in the *ELP1* transcript results in higher protein production (Fig. 6A, B, and E). The treatment resulted in a 2-fold increase in functional ELP1 protein in the brain and a 1.5-fold increase in the DRG (Fig. 6A and B). Because we used one retina for histology and the other to evaluate ELP splicing correction, there was no available tissue afterward to assess ELP1 protein expression. Together, these results provide the *in vivo* evidence that PTC258 increases the amount of ELP1 in CNS and PNS, thereby rescuing the primary neurologic FD phenotypes.

**Figure 6.**
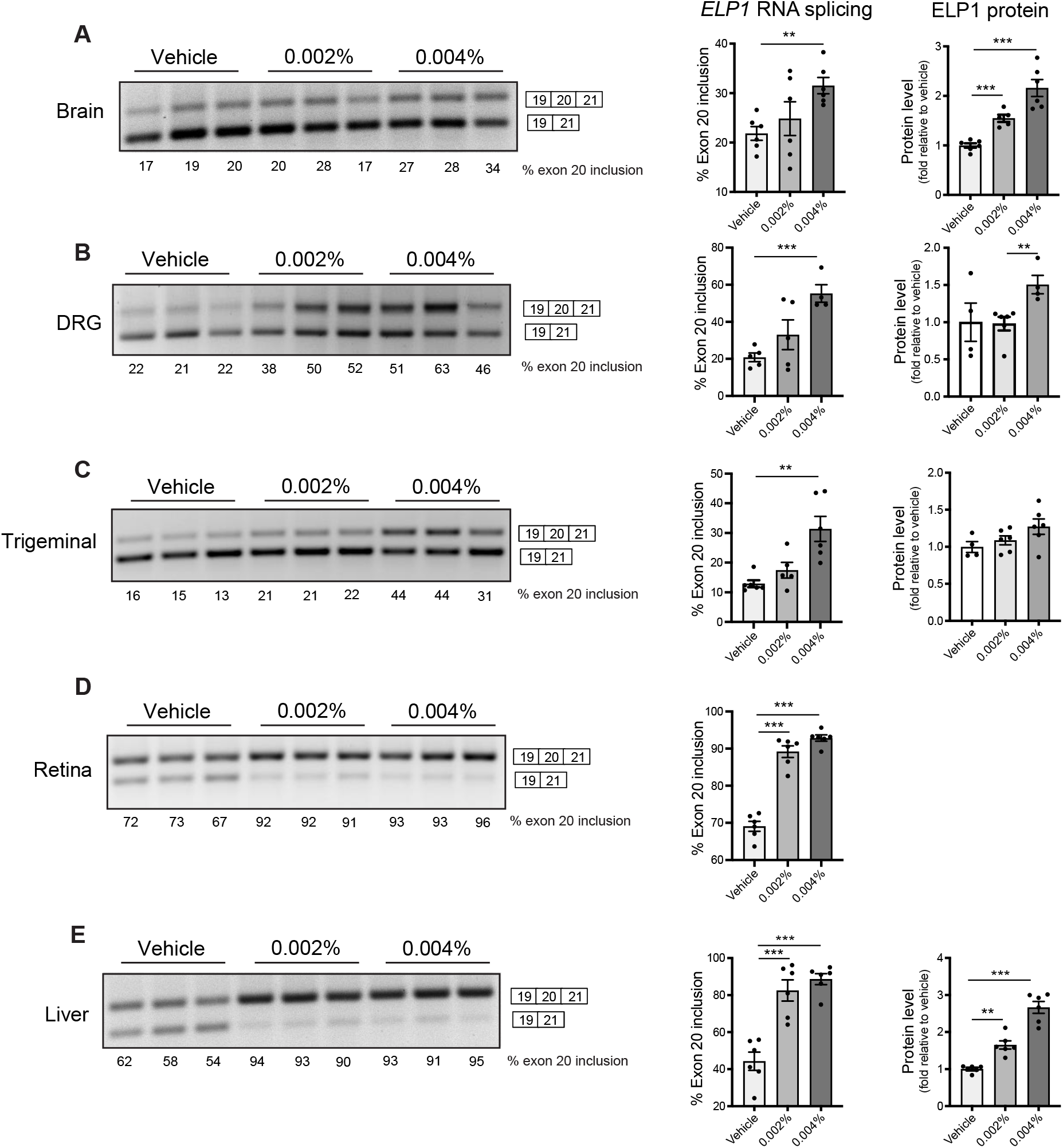
PTC258 treatment increases full-length *ELP1* transcript and protein in the FD mice. Representative splicing analysis of human *ELP1* transcripts (left), percent of exon 20 inclusion (middle) and levels of ELP1 (right) from vehicle-treated (n=5-6, light grey), 0.002% PTC258-treated (n=5-6, grey) and 0.004% PTC258-treated (n=4-6, dark grey) FD mice at 6 months of age. in brain (**A**), DRG (**B**), trigeminal (**C**), retina (**D**) and liver (**E**). The adjusted P values are displayed. **P< 0.01 and ***P< 0.001, two-tailed unpaired Student’s t-test with FDR correction. Data are shown as average ± s.e.m., each data point represents an individual animal.

## Discussion

FD is a complex sensory and autonomic neurodegenerative disease and, to date, there are no effective treatments to stop the continuous neuronal loss characteristic of this devastating disorder. A novel oral systemic therapy for FD will potentially revolutionize patient care. Importantly, because 99.5 % of FD patients are homozygous for the same splicing mutation in *ELP1*^72,73^, developing a treatment that precisely and efficiently targets the underlying genetic mechanism will benefit all patients. Many efforts have been undertaken to develop disease-modifying therapies including SMCs, ASO, and modified exon-specific U1 snRNAs ^54–56^. These modalities have shown promising effects in mice ^54,56,74^, confirming that *ELP1* splicing is a relevant therapeutic target. However, both ASO and U1 snRNA-based therapeutics have limitations, including poor brain penetration and cellular uptake, toxicity due to immune stimulation, and an invasive route of administration ^75–78^. The recent development of splicing targeted therapy for spinal muscular atrophy (SMA), another genetic disorder caused by a splicing alteration, has validated the utility of small molecules as a valuable therapeutic strategy for neurologic diseases ^79–83^. With the goal of developing an oral compound for FD, in this study, we have optimized the potency, efficacy, and distribution of SMCs that were initially generated as part of the NIH Blueprint Neurotherapeutics Network ^61^, and we have found a highly potent splicing modulator that efficiently passes the BBB and corrects *ELP1* splicing defect in PNS and CNS. We demonstrated that the novel compound, PTC258, efficiently restores correct *ELP1* splicing in several mouse tissues, including brain, and most importantly, prevents the progressive neuronal degeneration that is characteristic of FD.

Loss of proprioceptors accounts for many debilitating aspects of the disease, including progressive gait ataxia, spinal and craniofacial deformities, and respiratory insufficiency owing to neuromuscular incoordination^39,45^. We show that starting PTC258 treatment at birth rescues motor coordination in our FD phenotypic mouse model by preventing the loss of proprioceptors in the DRG. PTC258-treated FD mice showed increased DRG volume and number of proprioceptive neurons, indicating that increasing *ELP1* expression at birth is sufficient to prevent the loss of this subpopulation of neurons that play a critical role in disease progression. Similarly, we have tested the ability of our oral treatment to rescue progressive retinal degeneration. This is another significant debilitating aspect of the disease, as it severely affects patients’ quality of life. FD patients suffer from optic neuropathy featured by reductions of the RNFL due to progressive loss of the macular RGCs. They often become visually impaired or legally blind after their third decade of life ^35,36^. Postnatal retinal degeneration was also recapitulated in different mouse models of FD^59,84^. Supporting the idea that starting the treatment at birth can be sufficient to prevent this aspect of the disease. Oral administration of PTC258 significantly improved the thickness of RNFL and GCIPL by promoting RGC survival. To our knowledge, this is the first in vivo evidence of an oral treatment used to rescue retinal degeneration in a human genetic disease. We confirm that the phenotypic improvement promoted by PTC258 treatment correlates with a significant increase in full-length *ELP1* mRNA, which leads to an increase in full-length *ELP1* transcript and a 2-fold increase in functional ELP1 protein in the brain and 1.5-fold increase in the DRG.

Although we recognize that further safety and toxicity studies will be needed to move our SMCs to the clinic, we and others have shown that these compounds are very specific and selective in promoting the recruitment of the spliceosomal machinery at weakly defined 5’ splice sites ^9,85,86 60^. In fact, we have previously demonstrated that this class of compounds does not cause widespread changes in gene expression and splicing ^56,60^. Together, our work represents the first example of optimizing an orally bioavailable splicing modifier that corrects *ELP1* splicing in multiple tissues, including brain and retina, and provides the critical pre-clinical efficacy data needed for developing a novel oral treatment for FD patients.

## Materials and Methods

### Study Design

This study aimed to assess the therapeutic effectiveness of PTC258 in ameliorating neurological phenotypes *in vivo*. In this regard, we used the FD mouse model *TgFD9; Elp1^Δ20/flox^* because it recapitulates the same tissue-specific mis-splicing observed in individuals with FD and displays the hallmark symptoms of the disease, thus providing a powerful model for assessing the therapeutic efficacy of potential therapies. Treatment was started at birth to maximize the therapeutic value. At P0, regular mouse chow was replaced with vehicle diet (LabDiet^®^ 5P00) or PTC258 diets (LabDiet^®^ 5P00 w/ 20 ppm or 40 ppm PTC258), and the dam, randomly assigned, continued to be fed these diets until the time of weaning. At weaning, *TgFD9; Ikbkap^Δ20/flox^* mice were genotyped and maintained in the same treatment groups. We formulated PTC258 chow to provide each mouse with a dose of 3 mg/Kg/day or 6 mg/Kg/day. We have demonstrated that this dose was sufficient to significantly improve *ELP1* splicing and protein *in vivo* in the phenotypically normal mouse *TgFD9*.

All animal experiments were designed with a commitment to minimizing both the number of mice and their suffering. We designed our preclinical animal trial based on the published recommendations ^87^. To calculate appropriate sample sizes for the study, we performed a power analysis using the data generated from the previous kinetin efficacy study^56^. Thus, for statistical validity, we used at least n = 8 mice for phenotypic assessments, n = 5 mice for histological analysis of the DRG, and n = 4 to 6 mice for *ELP1* splicing and protein analysis. All analyses described in this study were conducted using animal samples from multiple litters; therefore, each unit (animal, cage, litter) represents a biological replicate. The numbers were not altered during the course of the study. The primary endpoints were predefined in advance based on our previous data ^56,59,64^. All data were included, and the criteria were established prospectively. Animals were assigned randomly to the vehicle- or PTC258-group using a randomization system devised by the MGH Biostatistics Center. The system consists of a box containing cards with either ‘vehicle diet’, ‘special diet 0.002%’ or ‘special diet 0.004%’ in random order, and animals were randomly assigned to the appropriate group by drawing a card. Investigators conducting the experiments were blind to genotype and treatment category.

### Animals

The generation of the *TgFD9* mouse line carrying the human *ELP1* transgene with the IVS20+6T>C mutation can be found in Hims et al. ^62^. Descriptions of the original targeting vector used to generate the *Elp1^flox^* allele and the strategy to generate the *Elp1^Δ20^* allele have been previously published ^88,89^. To generate the experimental *TgFD9; Elp1*^Δ*20/flox*^ mouse, we crossed the *TgFD9* transgenic mouse heterozygous for the *Elp1^flox^* allele (*TgFD9^+/−^*; *Elp1^flox/+^*) with each other. Pups were genotyped to identify those homozygotes for both the *TgFD9* transgene and the *Elp1^flox^* allele (F1: *TgFD9^+/+^; Elp1^flox/flox^*). These animals were then crossed with the mouse line heterozygous for the *Elp1*^Δ*20*^ allele (*E1p1*^Δ*20/+*^) to generate the FD mouse *TgFD9; Elp1^Δ20/flox^* (F2). Controls are littermates of the FD mice that carry the transgene but are phenotypically normal because they express the endogenous *Elp1 gene (TgFD9^+/−^; Elp1^+/+^, TgFD9^+/−^; Elp1 ^flox/+^ or TgFD9^+/−^; Elp1* ^Δ*20/+*^). The expected Mendelian ratio of *TgFD9; Elp1^Δ20/flox^* mouse using this breeding scheme was 1 in 2 (50%). However, since only about 60% of *TgFD9*; *Ikbkap^Δ20/flox^* mice survive postnatally ^64^, the actual ratio was 1:7 (28/184; 13%) for the vehicle-treated mice, 1:5 (28/125; 18.3%) for the 0.002% PTC258-treated mice and 1:4 (30/109; 21.6%) for the 0.004% PTC258-treated mice. Control and FD mice have a mixed background, including C57BL/6J and C57BL/6N. All the mice enrolled in the study were negative for the rd8 mutation.

The mice were housed in the animal facility at Massachusetts General Hospital (Boston, MA), provided with access to food and water ad libitum, and maintained on a 12-hour light/dark cycle. All experimental protocols were approved by the Institutional Animal Care and Use Committee of the Massachusetts General Hospital and were in accordance with NIH guidelines.

For routine genotyping of progeny, genomic DNA was prepared from tail biopsies, and PCR was carried out using the following primers - forward, 5`-TGATTGACACAGACTCTGGCCA-3’; reverse, 5`-CTTTCACTCTGAAATTACAGGAAG-3’- to discriminate the *Elp1* alleles and the primers - forward 5`-GCCATTGTACTGTTTGCGACT-3’; reverse, 5`-TGAGTGTCACGATTCTTTCTGC-3’- to detect the *TgFD9* transgene.

### Synthesis of PTC258 (2-[(2S)-2-Aminopropyl]-5-chloro-3-methyl-N-(2-thienylmethyl)thieno[3,2-b]pyridin-7-amine)

PTC258 was manufactured by PTC Therapeutic, Inc. All materials used in the studies were >99% pure, as assessed by analytical methods including NMR, HPLC and LC/MS.

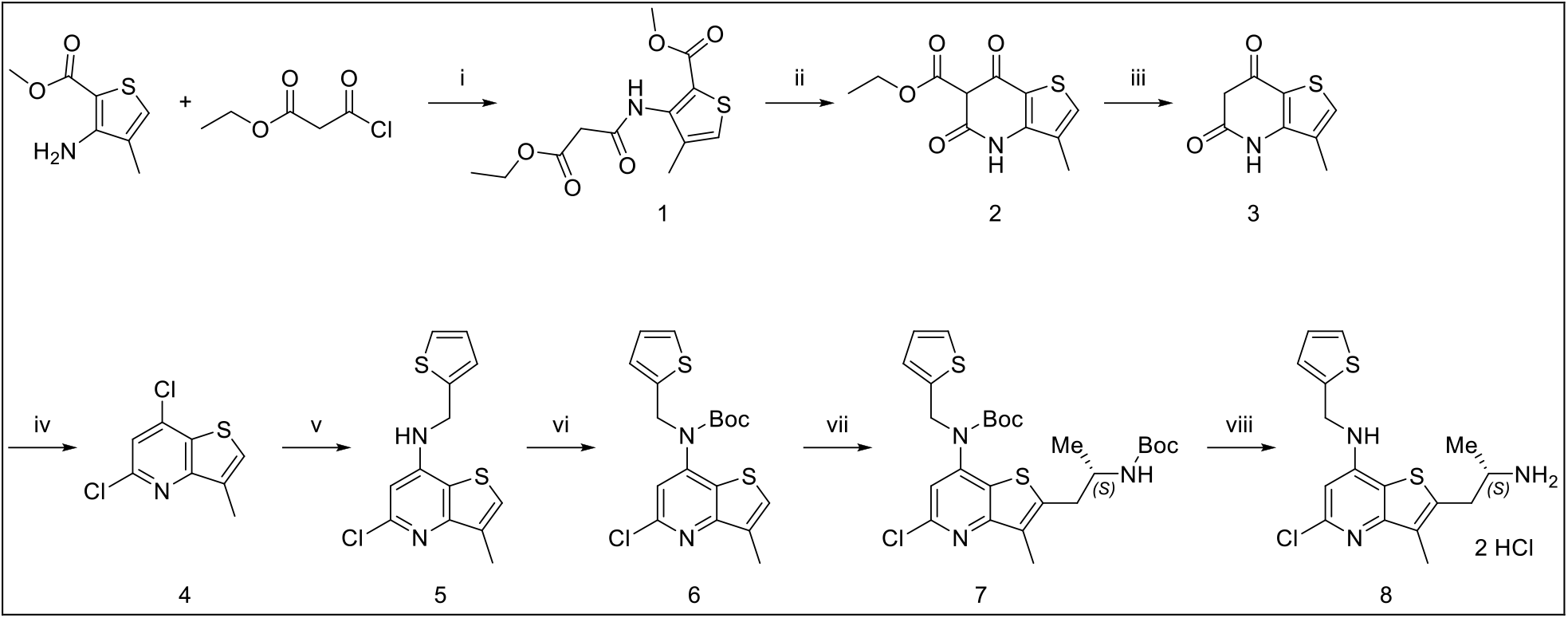

Reagents and conditions: (i) triethylamine, dichloromethane, 0°C to rt, 3 h, 98.4%; (ii) Na, EtOH, reflux, 12 h, 57.1%; (iii) NaOH, H2O, reflux, overnight, 75.3%; (iv) POCl3, N, N-dimethylaniline, reflux, overnight, 78%; (v) 2-thiophenemethylamine, DMSO, 100°C, 24 h, 42.7%; (vi) Boc2O, DMAP, DCM, rt, overnight, 74.8%; (vii) LDA, tert-butyl (4S)-4-methyl-2,2-dioxo-oxathiazolidine-3-carboxylate, −78°C, 20 min; then aq. citric acid, rt, 1 h, 78.4%; (viii) HCl in dioxane, rt, 15 min, 83%.

(1) Methyl 3-[(3-ethoxy-3-oxo-propanoyl)amino]-4-methyl-thiophene-2-carboxylate. To a solution of methyl 3-amino-4-methyl-thiophene-2-carboxylate (25.0 g, 146 mmol, 1.0 eq.) in dichloromethane (300 mL) was added triethylamine (40.7 mL, 292 mmol, 2.0 eq.). The mixture was cooled to 0°C and ethyl 3-chloro-3-oxo-propanoate (26.4 g, 175 mmol, 1.2 eq.) was slowly dropped in. After that, the mixture was stirred for 3 h at room temperature and quenched with brine (100 mL). The mixture was separated, and the aqueous phase was extracted with DCM (2 x 100 mL). The organic phase was dried over Na2SO4 and filtered. The filtrate was concentrated to give methyl 3-[(3-ethoxy-3-oxo-propanoyl)amino]-4-methyl-thiophene-2-carboxylate (41.0 g, 98.4% yield) as a brown oil, which was used in the next step without further purification. 1H NMR (chloroform-d) δ: 9.77 (br s, 1 H), 7.17 (d, J=0.8 Hz, 1 H), 4.31 (q, J=7.2 Hz, 2 H), 3.89 (s, 3H), 3.54 (s, 2H), 2.20 (d, J=0.8 Hz, 3H), 1.35 (t, J=7.2 Hz, 3 H).

(2) Ethyl 3-methyl-5,7-dioxo-4H-thieno[3,2-b]pyridine-6-carboxylate. At room temperature, sodium (1.65 g, 71.8 mmol, 0.5 eq.) was carefully dissolved in ethanol (150 mL) to give sodium ethanoate solution, which was mixed with methyl 3-[(3-ethoxy-3-oxo-propanoyl)amino]-4-methyl-thiophene-2-carboxylate (41.0 g, 144 mmol, 1.0 eq.). The mixture was refluxed for 12 h and then cooled to room temperature. Precipitation was formed and filtered. The filter cake was collected and dried in vacuo to give ethyl 3-methyl-5,7-dioxo-4H-thieno[3,2-b]pyridine-6-carboxylate (20.8 g, 57.1% yield) as a pale brown solid. MS m/z 254.1 [M+H]^+^; ^1^H NMR (D_2_O) δ: 7.16 (s, 1H), 4.18 (q, *J*=7.2 Hz, 2 H), 2.07 (s, 3H), 1.22 (t, *J*=7.2 Hz, 3H).

(3) 3-Methyl-4H-thieno[3,2-b]pyridine-5,7-dione. To a suspension of ethyl 3-methyl-5,7-dioxo-4H-thieno[3,2-b]pyridine-6-carboxylate (20.8 g, 82.1 mmol, 1.0 eq.) in water (200 mL) was added sodium hydroxide (5.97 g, 148 mmol, 1.8 eq.). The mixture was refluxed overnight, then cooled to room temperature. The pH was adjusted to 4~5 with conc HCl. Precipitation was formed and filtered. The filter cake was collected and dried in vacuo to give 3-methyl-4H-thieno[3,2-b]pyridine-5,7-dione (11.2 g, 75.3% yield) as a pale brown solid. MS m/z 182.1 [M+H]+; 1H NMR (DMSO-d) δ: 11.41 (br s, 2H), 7.45 (s, 1H), 5.62 (s, 1H), 2.24 (s, 3H).

(4) 5,7-Dichloro-3-methyl-thieno[3,2-b]pyridine. A mixture of 3-methyl-4H-thieno[3,2-b]pyridine-5,7-dione (5.0 g, 28 mmol, 1.0 eq.), POCl3 (60 mL), and N,N-dimethylaniline (1.5 mL, 12 mmol, 0.42 eq.) was refluxed overnight under N2 and then cooled. The mixture was concentrated via rotovap to remove most POCl3, then poured into ice water (20 mL). The precipitate was collected by filtration, washed twice with water, dried and purified by silica gel chromatography with dichloromethane to give 5,7-dichloro-3-methyl-thieno[3,2-b]pyridine (5.0 g, 78% yield) as a yellow solid. MS m/z 218.1, 220.0 [M+H]+; 1H NMR (chloroform-d) δ: 7.50 (d, J=0.8 Hz, 1 H), 7.37 (s, 1H), 2.52 (d, J=0.8 Hz, 3H).

(5) 5-Chloro-3-methyl-N-(2-thienylmethyl)thieno[3,2-b]pyridin-7-amine. A mixture of 5,7-dichloro-3-methyl-thieno[3,2-b]pyridine (1.0 g, 4.59 mmol, 1.00 eq.) and 2-thiophenemethylamine (5.2 g, 45.9 mmol, 10.0 eq.) in dimethyl sulfoxide (4.0 mL) was stirred at 100°C for 24 h, then cooled to room temperature, diluted with ethyl acetate and washed with water and brine, dried and evaporated. The residue was purified over silica with ethyl acetate and hexanes (5 to 35% gradient) to give 5-chloro-3-methyl-N-(2-thienylmethyl)thieno[3,2-b]pyridin-7-amine (0.58 g, 42.7% yield). MS m/z 295.1, 297.1 [M+H]+; 1H NMR (chloroform-d) δ: 7.32-7.30 (m, 1H), 7.26 (d, J=1.2 Hz, 1 H), 7.11-7.10 (m, 1H), 7.05-7.03 (m, 1H), 6.56 (s, 1H), 4.74-4.69 (m, 3H), 2.49 (s, 3H).

(6) tert-Butyl N-(5-chloro-3-methyl-thieno[3,2-b]pyridin-7-yl)-N-(2-thienylmethyl)carbamate. To a solution of 5-chloro-3-methyl-N-(2-thienylmethyl)thieno[3,2-b]pyridin-7-amine (0.58 g, 1.96 mmol, 1.0 eq.) and Boc2O (0.86 g, 3.92 mmol, 2.0 eq.) in dichloromethane (8.0 mL) was added DMAP (0.24 g, 1.96 mmol, 1.0 eq.) portion wise. Upon completion the solution was diluted with ethyl acetate and washed with water and brine, dried with Na2SO4 and concentrated. The residue was purified by column chromatography with ethyl acetate and hexanes (5 to 35% gradient) to furnish tert-butyl N-(5-chloro-3-methyl-thieno[3,2-b]pyridin-7-yl)-N-(2-thienylmethyl)carbamate (0.58 g, 74.8% yield). MS m/z 395.1, 397.1 [M+H]+; 1H NMR (chloroform-d) δ: 7.41 (s, 1 H), 7.24 (dd, J=1.0 Hz, 1 H), 7.06 (s, 1 H), 6.89 (dd, J=1.0 Hz, 1 H), 6.81 (d, J=1.0 Hz, 1 H), 5.07 (s, 2 H), 2.51 (d, J=1.1 Hz, 3 H), 1.47 (s, 9 H).

(7) tert-Butyl N-[2-[(2S)-2-(tert-butoxycarbonylamino)propyl]-5-chloro-3-methyl-thieno[3,2-b]pyridin-7-yl]-N-(2-thienylmethyl)carbamate. To a solution of tert-butyl N-(5-chloro-3-methyl-thieno[3,2-b]pyridin-7-yl)-N-(2-thienylmethyl)carbamate (0.58 g, 1.46 mmol, 1.0 eq.) in THF (6.0 mL) at −78°C was added 2.0 M LDA in THF/heptane/ethylbenzene (0.88 mL, 1.76 mmol, 1.2 eq.) dropwise. After 15 min a solution of tert-butyl (4S)-4-methyl-2,2-dioxo-oxathiazolidine-3-carboxylate (0.42 g, 1.76 mmol, 1.2 eq.) in THF (8.0 mL) was added. The yellow mixture was stirred at −78°C for 20 min then quenched with 1.0 M citric acid, followed by stirring at rt for 1 hour. The mixture was diluted with ethyl acetate, washed with water and brine, dried over sodium sulfate and evaporated. The residue was purified over silica gel with ethyl acetate and hexanes (3 to 30% gradient) to give tert-butyl N-[2-[(2S)-2-(tert-butoxycarbonylamino)propyl]-5-chloro-3-methyl-thieno[3,2-b]pyridin-7-yl]-N-(2-thienylmethyl)carbamate (0.63 g, 78.4% yield). MS m/z 553.0, 555.0 [M+H]+, 1H NMR (chloroform-d) δ: 7.22 (d, J=5.3 Hz, 1 H), 6.99 (s, 1 H), 6.86 (dd, J=5.0, 3.5 Hz, 1 H), 6.78 (d, J=2.9 Hz, 1 H), 5.02 (dd, J=1.0 Hz, 2 H), 4.46 (br s, 1 H), 4.01 (br s, 1 H), 3.07 - 3.16 (m, 1 H), 2.93 - 3.07 (m, 1 H), 2.41 (s, 3 H), 1.45 (s, 18 H), 1.14 (d, J=6.7 Hz, 3 H).

(8) 2-[(2S)-2-Aminopropyl]-5-chloro-3-methyl-N-(2-thienylmethyl)thieno[3,2-b]pyridin-7-amine, dihydrochloride. To a reaction tube with tert-butyl N-[2-[(2S)-2-(tert-butoxycarbonylamino)propyl]-5-chloro-3-methyl-thieno[3,2-b]pyridin-7-yl]-N-(2-thienylmethyl)carbamate (0.63 g, 1.15 mmol) was added hydrochloric acid (4.0 M) in dioxane (6.0 mL). After 15 minutes white precipitate appeared and UPLC confirmed the reaction was complete. To the mixture was added diethyl ether (15.0 mL). The precipitate was filtered. The filter cake was collected and dried under vacuum to furnish 2-[(2S)-2-aminopropyl]-5-chloro-3-methyl-N-(2-thienylmethyl)thieno[3,2-b]pyridin-7-amine, dihydrochloride (0.40 g, 83% yield). MS m/z 352.1, 354.1 [M+H]+, 1H NMR (MeOH-d4) δ: 7.38 (dd, J=5.2, 1.1 Hz, 1 H), 7.18 (d, J=2.8 Hz, 1 H), 7.10 (s, 1 H), 7.01 (dd, J=5.0, 3.5 Hz, 1 H), 4.97 (s, 2 H), 3.60 - 3.72 (m, 1 H), 3.37 - 3.44 (m, 1 H), 3.24 - 3.30 (m, 1 H), 2.47 (s, 3 H), 1.39 (d, J=6.6 Hz, 3 H).

### Catwalk analysis in mice

The Catwalk is an automated gait analysis system used to assess motor function and coordination in rodents. In brief, animals were allowed to walk on a green-illuminated glass platform contrasted by a red-illuminated ceiling, to allow for momentarily highlight of the footprints. A high-speed camera under the platform recorded movement and transferred the data to a computer, where paw prints were analyzed with the software CatWalk XT 10.6 (Noldus Inc., The Netherlands). During the data acquisition, each mouse was placed on the walkway in a dark environment and could walk freely in both directions with a minimum level of external disturbing factors. Experimental sessions typically lasted for 15-25 min. After data acquisition, each mouse was returned to its own home-cage, in order to reduce habituation of the animals to the environment and the appearance of unwanted behaviors (e.g. sniffing, rearing and sitting). Each mouse was tested on two consecutive days. Before the testing, the mice were allowed to acclimate to the experimental room for at least thirty minutes. Each experimental session lasted until 5 compliant runs were achieved. Compliance was defined as less than 60% variation in speed throughout the run with a minimum run duration of 0.5 seconds and a maximum run duration of 5 seconds. Any mouse having not completed five compliant runs in 25 minutes was removed from the apparatus and returned to the cage. The total compliant runs recorded from vehicle-treated control mice (n=20), vehicle-treated FD mice (n=16), 0.002% PTC258-treated FD (n=16) mice and 0.004% PTC258-treated FD mice (n=9) were respectively 32, 26, 25, and 15. The paw positions were first automatically labeled by the CatWalk system, then revised by an experienced observer. The minimum green intensity threshold was set at 0.10, the red ceiling light set at 17.7, the green walkway light set at 16.5, and the camera gain was set at 20.

### Spectral-Domain Optical coherence tomography (SD-OCT)

For *in vivo* imaging of the retina, mice were anesthetized by placing them in a mobile isoflurane induction chamber, and the vaporizer was set to an isoflurane concentration of 2% at 2 L/min O_2_. The mice’s pupils were dilated using 2.5% phenylephrine and 1% tropicamide. 0.5% proparacaine was used as a topical anesthetic during the procedure. SD-OCT imaging was performed using a Leica EnvisuR2210 OCT machine. Measurements were made at 100 μm from the optic nerve for the central retina and 1.5 mm from the optic nerve for the peripheral retina. Control and FD mice were analyzed at 3, 6, 12, and 18 months of age. Linear B-scans of the central and peripheral retina were performed, and the thickness of the RNFL and GCIPL layers were manually measured using Bioptigen InVivoView Clinc software. Each OCT image comprises 100 B-scans, with each B-scan consisting of 1000 A-scans. We then analyzed four representative images per mouse, two for each eye, and included measurements from both eyes.

### DRG immunohistochemistry

After euthanasia, L3 DRGs were dissected and fixed in 4% paraformaldehyde (PFA) overnight; afterward, the DRGs were washed for 24 h in PBS. The DRGs were then incubated in 30% sucrose, mounted in OCT compound, and stored at −80°C. 16 μm serial cryosections that spanned the whole ganglia were performed. Proprioceptive neurons were labeled with parvalbumin (PV; Synaptic System, guinea pig, 1:2000), and whole sensory neurons were labeled with fluorescent NueN staining (NueN; Chemicon International, mouse 1:500). We calculated the volume of the DRG by using ImageJ to measure, in every section, the area that was occupied by neuronal cell bodies and then multiplying the area of each section by its thickness (16 μm) to find the section volume. The sum of all the section volumes provided the DRG volume, expressed in mm^3 62^. We counted the number of total proprioceptive neurons per DRG by counting the number of proprioceptive neurons in every other section and then multiplying the average by the number of sections of each DRG.

### Retinal whole mounting and RGC counting

Staining and RGC counting of retinal whole mounts was performed according to the method previously described by Ueki et al. ^59,84,90,91^. Briefly, fixation of the eyes was performed at room temperature for 1 hour in 4% PFA, and the eyes were marked with a yellow tissue marking dye on the temporal surface. After fixation, retinae were isolated, with each temporal retina marked with a small cut. Relaxing cuts in the spherical retina were made on all four corners. Nonspecific binding was blocked by incubating with an animal-free blocker containing 0.5% Triton X-100 overnight at 4°C, and an anti-RPBMS antibody was applied overnight at 4°C. Retinae were incubated with secondary antibodies for 1 hour at room temperature and mounted on slides. Images were acquired using a LeicaDMi8 epifluorescent microscope and MetaMorph 4.2 acquisition software (Molecular Devices, San Jose, CA). Whole scans of complete flat-mount samples were obtained at 20X magnification using scan stage and autofocus. The total retinal area across the entire scan was approximately 14 mm^2^. With ImageJ software, the number of RPBMS+ cells were measured at 1×1 mm square at 1 mm from the ONH at superior, inferior, temporal, and nasal hemispheres ^59,84^. If a specific square area was damaged due to rips or folds in the retina, we have counted the RGCs in an adjacent undamaged area. Moreover, we have intentionally avoided counting RGCs in the edges of the retina because these areas usually have higher cell counts due to the downward pressure caused by the flattening cuts to the retina. We analyzed approximately 30% of the retina from one eye of each mouse. The investigator conducting the analysis was blinded to the genotype.

### RNA isolation and RT-PCR analysis of full-length and mutant *ELP1* transcripts in mouse tissues

Mice were euthanized, and brain, DRG, liver, lung, kidney, and heart tissues were removed and snap-frozen in liquid nitrogen. Tissues were homogenized in ice-cold TRI reagent (Molecular Research Center, Inc., Cincinnati, OH, USA), using a TissueLyser (Qiagen). Total RNA was extracted using the TRI reagent procedure provided by the manufacturer. The yield, purity, and quality of the total RNA for each sample were determined using a Nanodrop ND-1000 spectrophotometer. According to the manufacturer’s protocol, reverse transcription was performed using 1 μg of total RNA, Random Primers (Promega), and Superscript III reverse transcriptase (Invitrogen).

RT-PCR was performed using the cDNA equivalent of 100 ng of starting RNA in a 30-μl reaction, using GoTaq® green master mix (Promega) and 30 amplification cycles (94°C for 30 s, 58°C for 30 s, 72°C for 30 s). Human-specific *ELP1* primers - forward, 5`-CCTGAGCAG CAATCATGTG −3; reverse, 5`-TACATGGTCTTCGTGACATC-3’- were used to amplify human *ELP1* isoforms expressed from the transgene. PCR products were separated on 1.5% agarose gels and stained with ethidium bromide. The relative amounts of WT and mutant (Δ20) *ELP1* spliced isoforms in a single PCR were determined using ImageJ and the integrated density value for each band as previously described ^62,63^. The relative proportion of the WT isoform detected in a sample was calculated as a percentage.

### RT-qPCR analysis of full-length and mutant *ELP1* transcripts in mouse tissues

Mice were euthanized and brain, liver, lung, kidney, heart and skin tissues were removed and snap frozen in liquid nitrogen. Tissues were homogenized in ice-cold QIAzol Lysis Reagent (Qiagen), using Qiagen TissueLyser II (Qiagen). Total RNA was extracted using the QIAzol reagent procedure provided by the manufacturer. The yield, purity and quality of the total RNA for each sample were determined using a Nanodrop ND-1000 spectrophotometer. Full-length and mutant *ELP1* mRNA expression was quantified by quantitative real-time PCR (RT-qPCR) analysis using CFX384 Touch Real-Time PCR Detection System (BioRad). Reverse transcription and qPCR were carried out using One Step RT-qPCR (BioRad) according to the manufacturer’s recommendations. The mRNA levels of full-length *ELP1*, mutant Δ20 *ELP1* and *GAPDH* were quantified using Taqman-based RT-qPCR with a cDNA equivalent of 25 ng of starting RNA in a 20-μl reaction. To amplify the full-length *ELP1* isoform, FL *ELP1* primers forward, 5`- GAGCCCTGGTTTTAGCTCAG −3`; reverse, 5`- CATGCATTCAAATGCCTCTTT −3` and FL *ELP1* probe 5`- TCGGAAGTGGTTGGACAAACTTATGTTT-3` were used. To amplify the mutant (Δ20) *ELP1* spliced isoforms, Δ20 *ELP1* primers forward, 5`- CACAAAGCTTGTATTACAGACT −3`; reverse, 5`- GAAGGTTTCCACATTTCCAAG −3` and Δ20 *ELP1* probe 5`- CTCAATCTGATTTATGATCATAACCCTAAGGTG −3` were used to amplify the mutant (Δ20) *ELP1* spliced isoforms. The *ELP1* forward and reverse primers were each used at a final concentration of 0.4 μM. The *ELP1* probes were used at a final concentration of 0.15 μM. Mouse *GAPDH* mRNA was amplified using 20X gene expression PCR assay (Life Technologies, Inc.). RT-qPCR was carried out at the following temperatures for indicated times: Step 1: 48°C (15 min); Step 2: 95°C (15 min); Step 3: 95°C (15 sec); Step 4: 60°C (1 min); Steps 3 and 4 were repeated for 39 cycles. The Ct values for each mRNA were converted to mRNA abundance using actual PCR efficiencies. *ELP1* FL and Δ20 mRNAs were normalized to *GAPDH* and vehicle controls and plotted as fold change compared to vehicle treatment. Data were analyzed using the SDS software.

### ELP1 protein quantification in human fibroblasts using Homogeneous Time Resolved Fluorescence (HTRF) assay

GM04589 patient fibroblasts were thawed and incubated in Dulbecco’s Modified Eagle Medium (DMEM-10%) fetal bovine (FBS) for 72 hours. Cells were trypsinized, counted, and resuspended to a concentration of 50,000 cells/ml in DMEM-10% FBS. Aliquots (199 μl) of the cell suspensions were plated at 10,000 cells per well in a 96-well microtiter plate and incubated for 3 to 5 hours. Two sets of control wells were included in each plate, 6 wells received DMSO at a final concentration of 0.5% and 6 wells were filled with cell culture medium without cells and served as blank wells. PTC258 was serially diluted 3.16-fold (i.e., half log-10 dilution scheme) in 100% DMSO to generate a 7-point concentration curve. A 1 μl aliquot of 200x compound solution was transferred to cell-containing wells, and cells were incubated for 48 hours in a cell culture incubator (37°C, 5% CO2, 100% relative humidity). Three independent samples were set up for each compound concentration. After 48 hours, the supernatant was removed from the cells and 50 μl of the 1x LB4 (lysis buffer), containing protease inhibitors, was added to the cells and incubated with shaking at room temperature for 1 hour. A 36 μl aliquot of this lysate was subsequently transferred to a 384 well plate containing 4 μl of the antibody solution (1:50 dilution of anti-ELP1 d2(CisBio) and anti-ELP1 K(CisBio) in detection buffer). The 384-well plate was then centrifuged for 1 minute to bring the solutions to the bottom of the plate and incubated overnight at 4°C. Fluorescence emission for each well of the plate at 665 nm (acceptor) and 620 nm (donor) was measured on the EnVision plate reader (Perkin Elmer).

### Meso Scale Discovery (MSD) immunoassay for ELP1 protein quantification in mouse tissues

Tissue samples were collected in Safe-Lock tubes (Eppendorf), snap-frozen in liquid nitrogen, weighed and homogenized on the TissueLyzer II (Qiagen) in RIPA buffer (Tris-HCl 50 mM, pH 7.4; NaCl 150 mM; NP-40 1%; sodium deoxycholate 0.5%; SDS 0.1%) containing a cocktail of protease inhibitors (Roche) at a tissue-weight to RIPA buffer volume of 50 mg/ml. The samples were then centrifuged for 20 min at 14,000 x g in a microcentrifuge. The homogenates were transferred to a 96-well plate and were diluted in RIPA buffer to ~1 mg/ml for ELP1-MSD and ~ 0.5 mg/mL for total protein measurement using the BCA protein assay (Pierce). Samples were run in duplicate and averaged. The MSD sandwich immunoassay was performed according to the manufacturer’s (Meso Scale Diagnostics) protocol. 25 μl of the diluted tissue homogenates were transferred to a 96-well standard MSD plate coated with 0.5 μg/ml of capture antibody, rabbit monoclonal anti-ELP1 antibody from Abcam #ab179437, in PBS and incubated overnight at 4°C. The primary detection antibody, mouse anti-IKAP (33) from Santa Cruz Biotechnology #SC-136412, was used at 0.5 μg/ml and incubated for 2-3 hours at room temperature. Sulfo-Tag antibody, Goat anti-mouse from MSD #R32AC-1, was used at 0.5 μg/ml and incubated for 1 hour at room temperature. Sector Imager S600 (Meso Scale Diagnostics) was used to read the plates. The level of ELP1 in the tissues from kinetin-treated mice was normalized to the ELP1 level in the control tissues and plotted as fold change over controls.

### Statistical analysis

We performed an unpaired Student’s t-test using GraphPad Prism 7 software to determine the statistical differences between two groups. Every time one group was compared to more than another group, we applied the false discovery rate (FDR) correction and reported the FDR-adjusted *P* values. Every time different treatment groups were compared to the same control group, we applied one-way ANOVA with Dunnett’s multiple comparison test and reported the adjusted *P* values. For all experiments, a criterion α level was set at 0.05.

## Conflict of Interest

The authors declare competing financial interests.

Jana Narasimhan, Vijayalakshmi Gabbeta, Shivani Grover, Amal Dakka, Anna Mollin, Stephen Jung, Xin Zhao, Nanjing Zhang, Sophie Zhang, Michael Arnold, Matthew G. Woll, Nikolai A. Naryshkin, Marla Weetall are/were employees of PTC Therapeutics, Inc., a biotechnology company. In connection with such employment, the authors receive salary, benefits and stock-based compensation, including stock options, restricted stock, other stock-related grants, and the right to purchase discounted stock through PTC’s employee stock purchase plan.

Funding: Research support from PTC Therapeutics, Inc. (S.A.S. and E.M.).

Personal financial interests: Susan A. Slaugenhaupt is a paid consultant to PTC Therapeutics and is an inventor on several U.S. and foreign patents and patent applications assigned to the Massachusetts General Hospital, including U.S Patents 8,729,025 and 9,265,766, both entitled “Methods for altering mRNA splicing and treating familial dysautonomia by administering benzyladenine,” filed on August 31, 2012 and May 19, 2014 and related to use of kinetin; and U.S. Patent 10,675,475 entitled, “Compounds for improving mRNA splicing” filed on July 14, 2017 and related to use of BPN-15477.

Elisabetta Morini, Vijayalakshmi Gabbeta, Amal Dakka, Nikolai A. Naryshkin, and Susan A. Slaugenhaupt are inventors on an International Patent Application Number PCT/US2021/012103, assigned to Massachusetts General Hospital and PTC Therapeutics entitled “RNA Splicing Modulation” related to use of BPN-15477 in modulating splicing.

All other authors declare no competing interests.

## Acknowledgments

We thank Dr. Horacio Kaufmann at the Dysautonomia Treatment and Evaluation Center at New York University Medical School and Dr. Frances Lefcort at Montana State University for their long-standing collaboration and helpful discussions. We are also grateful to Drs. Ioannis Dragatsis and Paula Dietrich at the University of Tennessee for the gift of the *Elp1^flox/+^* and *Elp1^Δ20/+^* mice and their collaborative effort to generate the FD phenotypic mouse model. This work was supported by National Institutes of Health (NIH) grants (R37NS095640 to S.A.S.).

