## Supplementary Materials for "Development of a novel oral treatment that rescues gait ataxia and retinal degeneration in a phenotypic mouse model of familial dysautonomia"

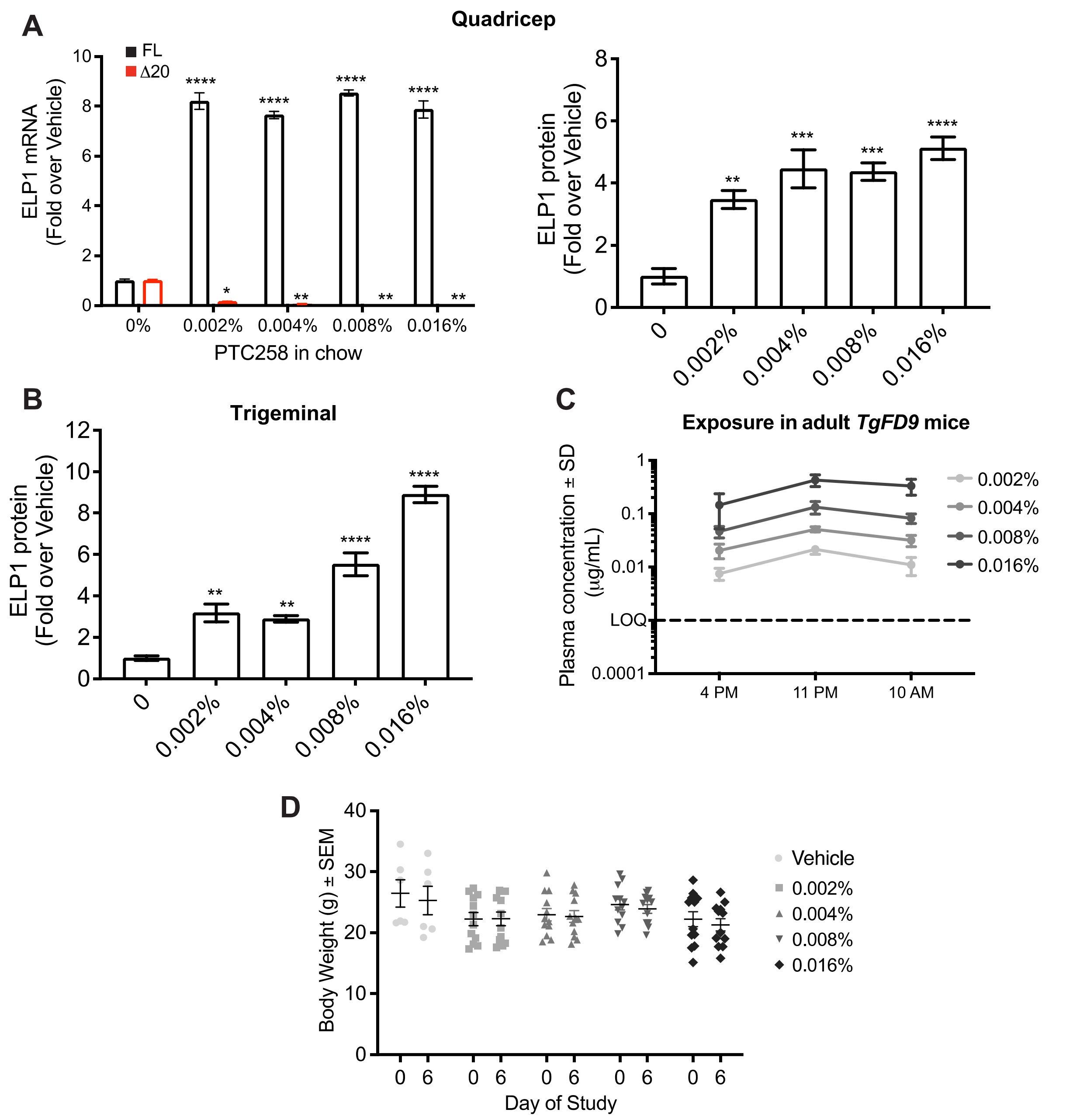


**Supplementary Figure 1. Oral administration of PTC258 to the *TgFD9* transgenic mouse.** (A and B) Relative expression of full-length (FL) and Δ20 ELP1 mRNA (left panel), and ELP1 protein quantification (right panel) in quadricep and trigeminal after oral administration of PTC258 in chow ranging from 0.002% to 0.016% in adult transgenic *TgFD9* mouse (n =6-11). (C) Plasma concentration of PTC258 at different timepoints: 4 pm, 11 pm and 10 am. The levels of compound were measured using mass spectrometry. (D) Weight assessment of *TgFD9* mice in different treatment groups. The adjusted P values are displayed. *P< 0.05, **P< 0.01, ***P< 0.001 and ****P< 0.0001, one-way ANOVA with Dunnett's multiple comparison test. Data are shown as average ± s.e.m.

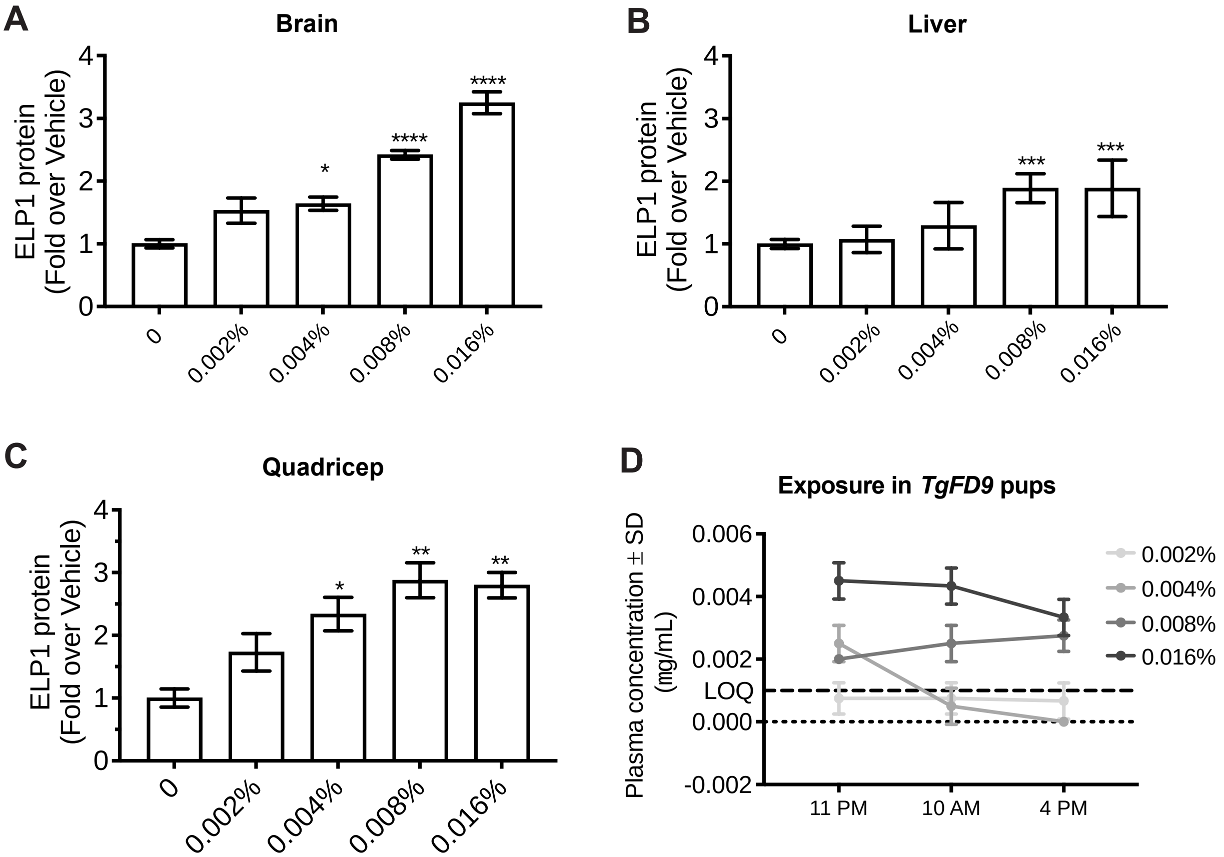


**Supplementary Figure 2. PTC258 improves ELP1 protein in nursing pups.** At the day of delivery (P0) the dams were randomly assigned to vehicle diet or PTC258 diet, and, continued to be fed these diets until the time of sacrifice (P6). ELP1 protein was analyzed in nursing pups carrying the human transgene with the major FD splicing mutation, *TgFD9*. (A-C) ELP1 protein quantification in brain (A), liver (B), and quadricep (C) from *TgFD9* pups (n= 3 in the vehicle-treated group, n=6-12 in the PTC258-treated groups) treated with special formulated chow ranging from 0.002% to 0.016%. (D) Plasma concentration of PTC258 at different timepoints: 11 pm, 10 am, and 4 pm. The levels of compound were measured using mass spectrometry. The adjusted P values are displayed. *P< 0.05, **P< 0.01, ***P< 0.001 and ****P< 0.0001, one-way ANOVA one-way ANOVA with Dunnett's multiple comparison test. Data are shown as average ± s.e.m.

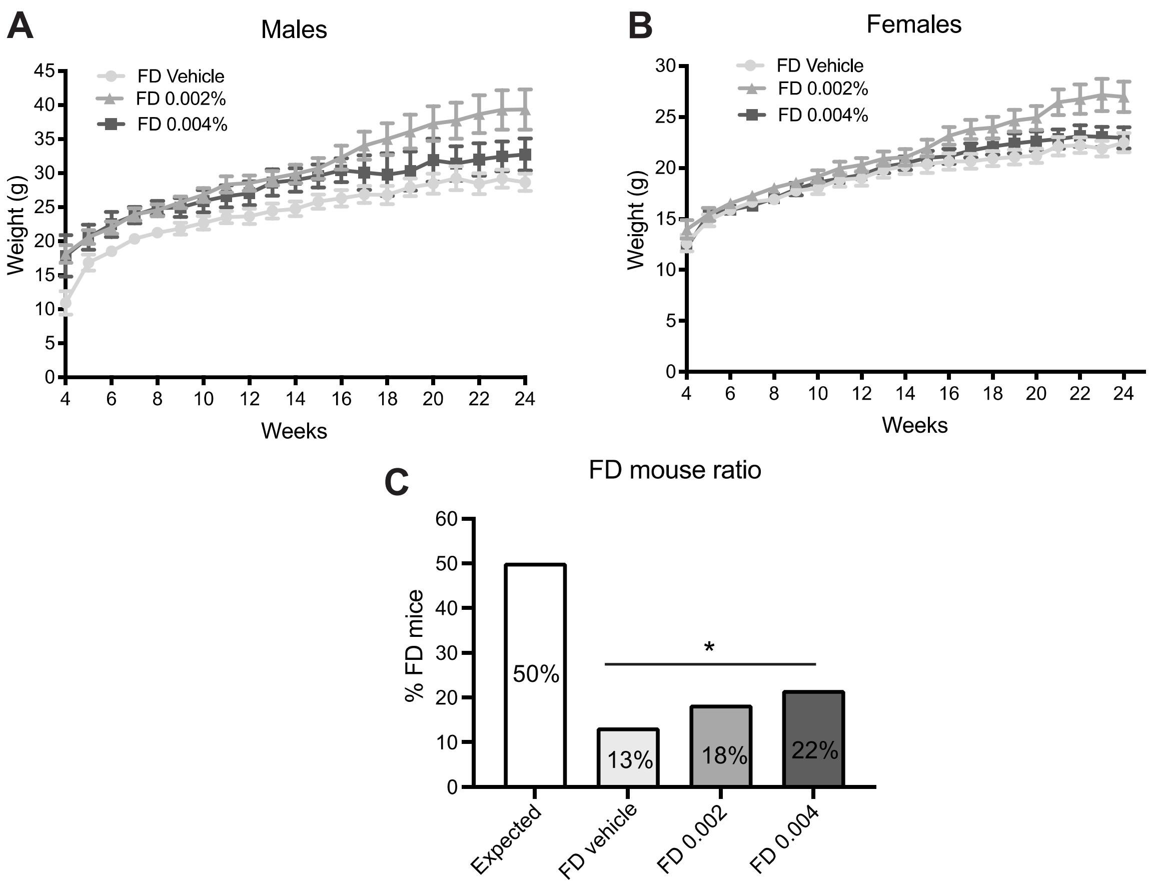


**Supplementary Figure 3. PTC258 treatment was well tolerated and increased the ratio of the FD mice.** (A) Postnatal growth curves for vehicle-treated (n=9, light gray), 0.002% PTC258-treated (n=7, gray) and 0.004% PTC258-treated (n=4, dark gray) FD male mice. (B) Postnatal growth curves for vehicle-treated (n=9, light gray), 0.002% PTC258-treated (n=15, gray) and 0.004% PTC258-treated (n=9, dark gray) FD female mice. Means and s.e.m. are shown. (C) The expected Mendelian ratio of $TgFD9^{+/-}$; $Elp1^{\Delta 20/flox}$ mice obtained by crossing $TgFD9^{+/+}$; $Elp1^{flox/flox}$ mice × $Elp1^{\Delta 20/+}$ mice is 50% (white bar). The actual ratio was 13% (28/184) for the vehicle-treated mice, 18.3% (28/125) for the 0.002% PTC258-treated mice and 21.6% (30/109) for the 0.004% PTC258-treated mice. A significant difference was observed in the actual ratio between vehicle-treated and 0.004% PTC258-treated FD mice. *$P < 0.05$, $\chi^2$ –Test.

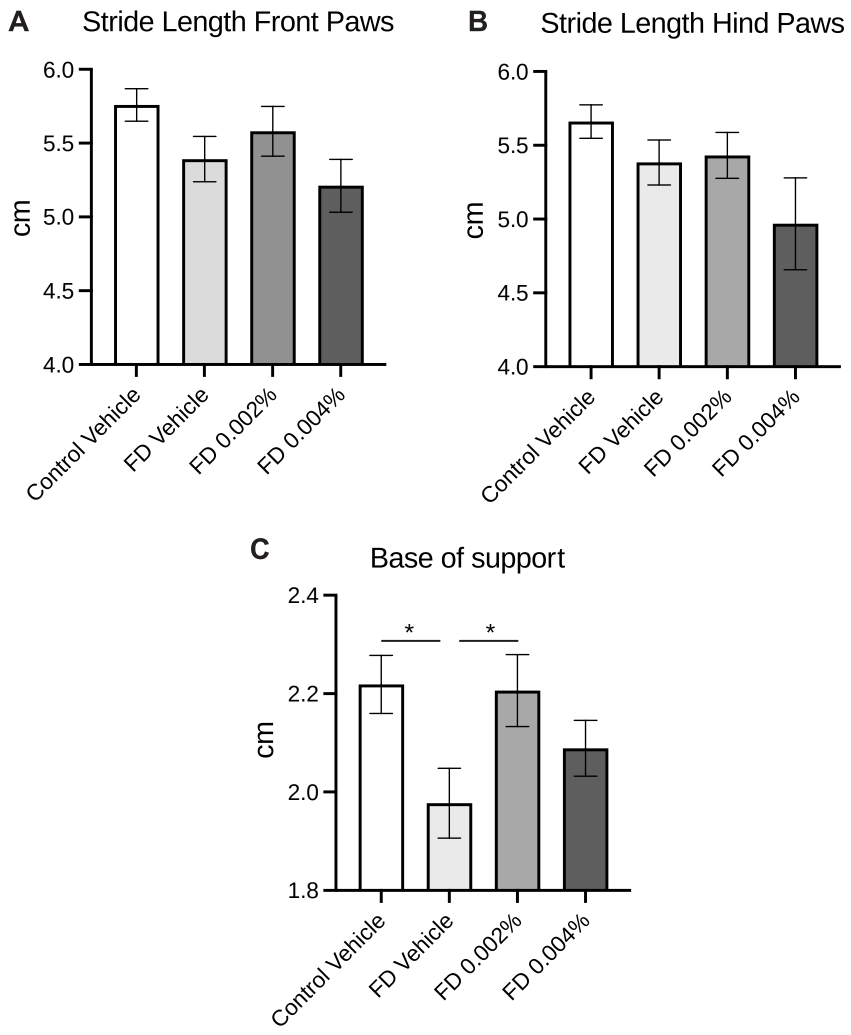


**Supplementary Figure 4. Catwalk analysis in 3 month-old FD mice treated with increasing doses of PTC-258.** Measurements of stride length front paws (A), stride length hind paws (B) and base of support hind paws (D) in vehicle-treated control mice (n=11) and vehicle-treated (n=10), 0.002% PTC258-treated (n=10) and 0.004% PTC258-treated (n=9) FD mice at 3 months of age. Stride length is defined as the distance between successive placements of the same paw (blue double-headed arrows), while the base of support represents the mean distance between hind paws (red double-headed arrows). The adjusted P values are displayed. *P< 0.05, two-tailed unpaired Student's t-test with FDR correction. Data are shown as average ± s.e.m.
